# The distribution of particle-associated *Prochlorococcus* across the global oceans

**DOI:** 10.64898/2026.08.24.746807

**Authors:** Maya I. Anjur-Dietrich, Nhi N. Vo, Katelyn G. Jones, James I. Mullet, Sierra M. Parker, Kurt G. Castro, Ashley M. Stein, Samantha M. Silvestri, Steven J. Biller, Krista Longnecker, Sallie W. Chisholm

## Abstract

The picocyanobacterium *Prochlorococcus* is a fundamental contributor to ocean primary productivity. While its free-living population has been extensively studied, primarily using flow cytometric analyses, the size and distribution of its particle-associated population is not well understood. Using filter fractionated samples from cruises in the Pacific Ocean, Atlantic Ocean, and Mediterranean Sea, we generated metagenomic data using internal standards, yielding absolute genome equivalent counts of *Prochlorococcus* cells in different size fractions. We used these data to model a relationship between relative and absolute genome equivalent counts, yielding a correction factor that we validated using published datasets. We then applied the correction factor to size-fractionated global metagenomic data from the TARA Oceans Project, which has widespread *Prochlorococcus* cells in size fractions >1.6 μm throughout the transects, to calculate the fraction of the total *Prochlorococcus* population in large size fractions. The “particle-associated” population fraction increased with net primary productivity. Dissolved inorganic carbon was also directly correlated with increased particle association, which, combined with other evidence, could indicate an association with upwelling. We also examined the relationship between particle-associated population and carbon export at 150 m by incorporating published estimates of carbon flux based on TARA optical scattering data. This study highlights the potential importance of particle-associated *Prochlorococcus* to carbon flux in marine ecosystems and offers a way to convert relative to absolute genome equivalents of microorganisms in archival metagenomic datasets.

## Introduction

Marine microbes, such as the picocyanobacterium *Prochlorococcus*, are major contributors to ocean biogeochemical cycles and primary productivity. *Prochlorococcus’*s distribution across the global oceans has primarily been studied in its free-living form ^1–4^. Beyond some evidence of *Prochlorococcus* genetic signatures on particles ^5–9^, little is known about the abundance and community structure of *Prochlorococcus* on marine particles. Resolving whether *Prochlorococcus* distributions differ between free-living and particle-associated populations has been hindered by limitations in both of the standard methods for detecting these cells—flow cytometry and metagenomics. In flow cytometric analysis, *Prochlorococcus* is identified as the smallest particle, as measured by forward light scatter, that emits laser-induced red fluorescence from the excitement of chlorophyll. Aggregated or particle-associated cells have higher scattering signals and are mixed with signals from larger eukaryotic phytoplankton and thus cannot be identified as *Prochlorococcus*. Metagenomics easily classifies *Prochlorococcus* genetic signatures and, when paired with size fractionation, offers an approach for differentiating between free-living and particle-associated cells. To compare between samples, however, requires absolute quantification of genome counts in the different fractions.

Typically, metagenomic surveys are reported as compositional datasets ^10^, or relative abundances based on a variety of normalization strategies ^11–14^. As these data only report proportions between types of microbes within a single sample, methods of whole-genome internal standards have been developed that allow for quantitative comparison of absolute abundances between samples ^15,16^. Datasets with internal standards also provide opportunities to explore how to estimate absolute abundances from relative metagenomic sampling, for instance by normalizing amplicon data using flow cytometry cell counts ^17^. Given the rich history of marine metagenomic sampling without internal standards, these types of correction methods provide an opportunity for quantitative reanalysis of vast existing datasets and are particularly useful for understanding the distribution of *Prochlorococcus*, which is not only found in free-living size fractions but is also involved in microbial communities on particles ^5–9^.

Here, we sequenced internal standard-normalized metagenomes from four ocean sampling expeditions to quantitatively investigate distributions of particle-associated *Prochlorococcus* populations (i.e. cells in size fractions >1.2 μm, as *Prochlorococcus* cells are ∼600 nm in diameter ^18^). Since this was a pilot study, we framed our inquiries to learn as much as possible while taking advantage of existing cruise samples which, in two cases, had already been collected and size-fractionated, meaning we had no input on filter sizes. As such, the size fractions vary between cruises. As we discuss below, more targeted sampling designed to separate free-living and particle-associated *Prochlorococcus* would enable more precise quantification. However, taking advantage of the available samples, we used these size-fractionated metagenomic data to determine a relationship between relative and absolute genome equivalent counts. We applied a correction factor derived from this analysis to data from the TARA Oceans Project ^19–21^ to estimate free-living and particle-associated *Prochlorococcus* population fractions in samples from around the globe. Finally, we examined patterns in *Prochlorococcus* particle association as a function of environmental conditions to form hypotheses regarding factors that might influence this feature of *Prochlorococcus* ecology.

## Results and Discussion

### Prochlorococcus populations are found in large size fractions

To estimate the fraction of *Prochlorococcus* associated with particles, we first obtained serially-filtered, size-fractionated samples from 3 ocean sampling expeditions: FK180310 (SCOPE Falkor Cruise) in 2018 at Station ALOHA, SoMMoS (Southeastern Mediterranean Monthly cruise Series) in 2018-2019 at the Texas-Haifa Eastern Mediterranean Observatory 2 ^22^, and BATS AE2412 in 2024 from the Bermuda Atlantic Time-series Study site (Fig. 1A, with size fractions described in Fig. 1B and Supp. Table 1). We extracted all samples with added internal standards and combined the normalized metagenomes with previously reported data ^5^ collected during HOT346 at Station ALOHA (see Methods). For readability, we refer to samples from FK180310 as “Falkor”, samples from BATS AE2142 as “BATS”, samples from SoMMoS as “SoMMoS”, and samples from HOT346 as “HOT”. We also define the “free-living” size fraction as 0.2-1.2 μm, and the “particle-associated” size fractions as any >1.2 μm. Finally, we note that for Falkor and SoMMoS, the smallest fraction is 0.22-5 μm, which includes both free-living cells and small particles and aggregates; in our discussion, we define this as the “small” size fraction. Below, we discuss how these samples can be used to determine the lower bound of the particle-associated *Prochlorococcus* fraction, and data from these cruises will be represented with squares throughout to indicate the lack of a true free-living fraction.

**Figure 1.**
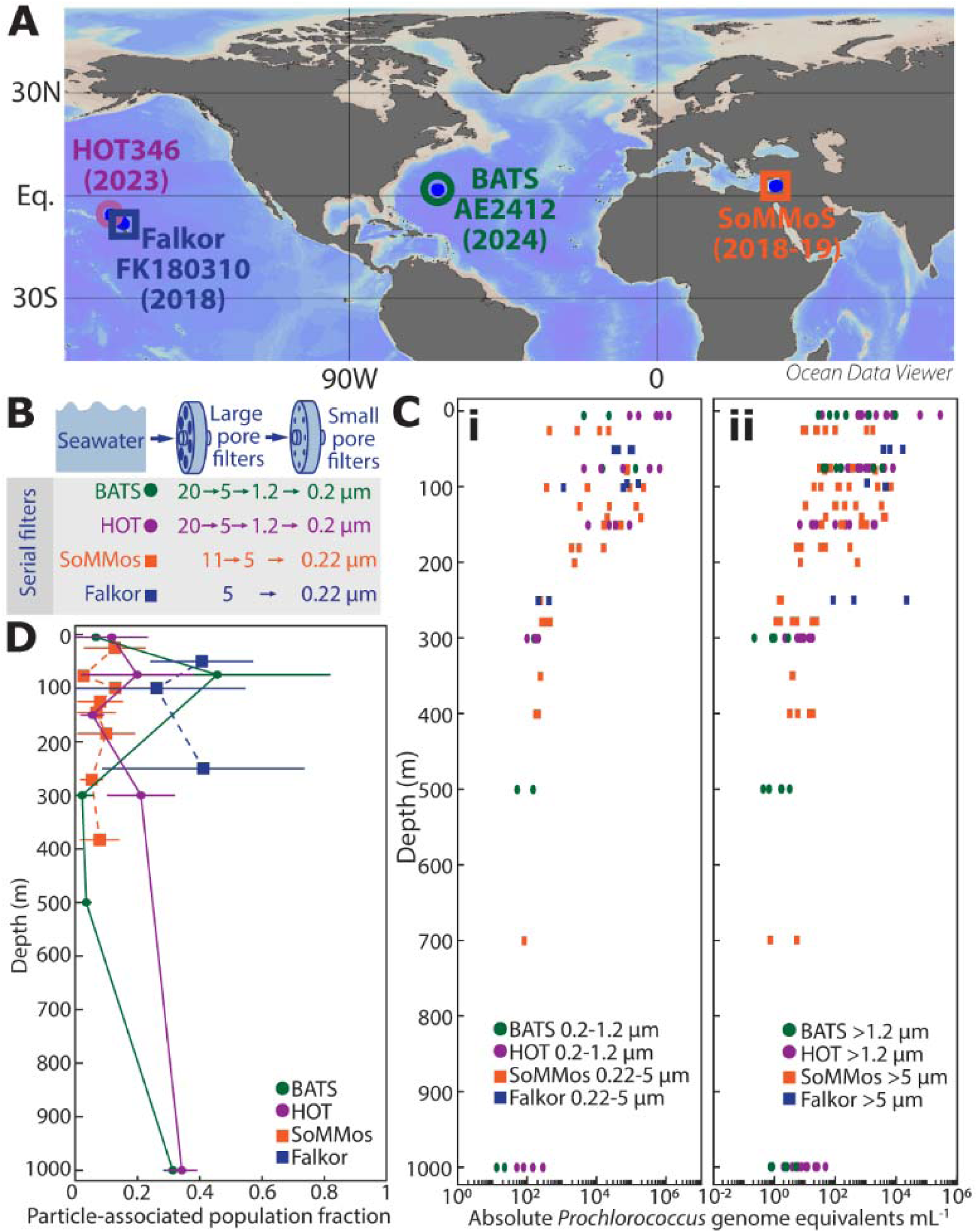
Fraction of *Prochlorococcus* genome equivalents found in various serial filtration size fractions using internal standard-normalized metagenomic samples. (A) Cruise coordinates and dates. (Cruises are abbreviated in the rest of the figure as follows: HOT346 as “HOT”, BATS AE2412 as “BATS”, FK180310 as “Falkor”, and SoMMoS as “SoMMoS”.) (B) Schematic of serial filtration for all cruises. Falkor and SoMMoS cruises will be represented with squares throughout to indicate that these sampling schemes did not have a true free-living fraction. (C) (i) Internal standard-normalized *Prochlorococcus* genome equivalent counts in the free-living fraction (0.2-1.2 μm) for HOT and BATS (circles) and the smallest size fraction (0.22-5 μm) for Falkor and SoMMoS (squares) as a function of depth. (ii) Internal standard-normalized *Prochlorococcus* genome equivalent counts in the particle-associated size fractions for HOT (1.2-5 μm, 5-20 μm, and >20 μm), BATS(1.2-5 μm, 5-20 μm, and >20 μm), SoMMoS (5-11 μm, >11 μm), and Falkor (>5 μm). (D) Fraction of total *Prochlorococcus* genome equivalent counts in the particle-associated fraction for HOT and BATS (>1.2 μm) (error bars SEM of samples at each depth per cruise), and the lower-bound fraction of total *Prochlorococcus* genome equivalent counts in the particle-associated fraction for Falkor and SoMMoS (>5 μm). (See also Supp. Fig. 1A-D)

We found expected and similar trends of free-living *Prochlorococcus* (genome equivalents mL^-1^ of seawater) in both the HOT and BATS 0.2-1.2 μm samples (Fig. 1Ci, dots), where free-living cells were most numerous in the upper euphotic zone with population sizes decreasing with depth. We also found similar trends in the SoMMoS and Falkor 0.22-5 μm samples (Fig. 1Ci, squares), even with the larger range of this size fraction, which is consistent with the majority of *Prochlorococcus* cells being free-living. Next, we looked at the genome equivalents in the particle-associated size fractions, either >1.2 μm for HOT and BATS samples or >5 μm for SoMMoS and Falkor samples. We found that the number of genome equivalents in the particle-associated size fractions (HOT, BATS: 1.2-5 μm, 5-20 μm, and >20 μm; SoMMoS: 5-11 μm, and >11 μm; Falkor: >5 μm) was about one to two orders of magnitude lower than in the small size fractions, and that the number also decreased with depth (Fig. 1Cii). In this case, we have shown each individual size fraction as a single datapoint, indicating a relatively consistent distribution of *Prochlorococcus* genomes across multiple size ranges between sampling sites.

We next described the particle-associated fraction of the total *Prochlorococcus* population, i.e. the sum of genome equivalents in all particle-associated size fractions divided by the sum of genome equivalents in all size fractions (see detailed calculations in Methods). For the HOT and BATS samples, particle-associated fractions increased around 75 m relative to the fraction at the surface (Fig 1D). These two cruises also included mesopelagic sampling, and there was an increase in the particle-associated fraction at 1000 m relative to the fraction at the surface, showing that particle association varies with depth across multiple oceans. For the Falkor and SoMMoS samples, we could only calculate a lower bound on the particle-associated population fraction: since some small particles (and thus any *Prochlorococcus* cells on those particles) were in the smallest size fraction, the number of genome equivalents in the larger size fractions would be an underestimate of the total particle-associated cells. For the Falkor samples, we found that the particle-associated fraction was on average higher than in the other cruises, which may relate to sampling a cyclonic eddy ^23^. SoMMoS samples had relatively low—but consistently measurable—particle-associated population fractions overall (Fig. 1D, orange).

The total *Prochlorococcus* population is broadly split into high light-adapted and low light-adapted ecotypes, each with differing genome content, optimal growth conditions, and spatial and depth distributions^1–3,24,25^. While the capacity for attachment is found across *Prochlorococcus* ecotypes, more cells from low light-adapted ecotypes are found on particles in the upper euphotic ^5^. Similarly, for all four datasets, the particle-associated fraction of low light-adapted ecotypes was greater than that of high light-adapted ecotypes at the surface (Supp. Fig. 1A-D). We also combined metagenomic analysis with cell counts to estimate the number of particle-associated *Prochlorococcus* cells (Supp. Fig. 1E, Supp. Table 2, see also Methods), although flow cytometry and metagenomic samples were not paired across the four cruises. Data from these cruises indicate that a significant fraction of *Prochlorococcus* cells is associated with particles across depths and in multiple different oceans.

### The relationship between genome size-normalized and internal standard-normalized counts

The DNA of the majority of oceanic metagenomic samples has been extracted without internal standards for normalization and yet, when samples were size-fractionated, they can hold valuable clues regarding the fraction of *Prochlorococcus* cells attached to particles. Since particle-associated population fractions are based on ratios of genome equivalents between size fractions, the ability to quantitatively relate counts between samples is crucial for utilizing publicly available data to address these questions. To this end, we first compared the particle-associated fractions calculated using either genome size- or internal standard-normalized counts (Fig. 2A) for both the “true” particle-associated fractions (for HOT and BATS) and the lower bound estimates (for Falkor and SoMMoS) (again, see detailed calculations in Methods). Genome size-normalized counts are determined by dividing the number of *Prochlorococcus* reads in a metagenomic sample by the average total length of a set of core genes in either high light-adapted or low light-adapted ecotypes. Internal standard-normalized counts add another layer of normalization by adjusting for the extraction efficiency based on a known DNA standard (see Methods for details). Essentially, genome size-normalized counts allow for comparison between microbial population counts within a sample, while internal standard-normalized counts can be quantitatively compared between samples. Particle-associated fractions calculated using genome size-normalized counts were in general much larger than the fractions calculated from internal standard-normalized counts, regardless of whether the smallest size fraction truly captured only free-living cells, showing that genome size-normalized counts lead to overestimated particle-associated populations (Fig. 2A). Therefore, genome size-normalized counts—the typical genome equivalent count reported in metagenomic samples—cannot be used to quantitatively compare samples or accurately measure population fractions split between free-living and particle-associated states, even with optimal sample fractionation.

**Figure 2.**
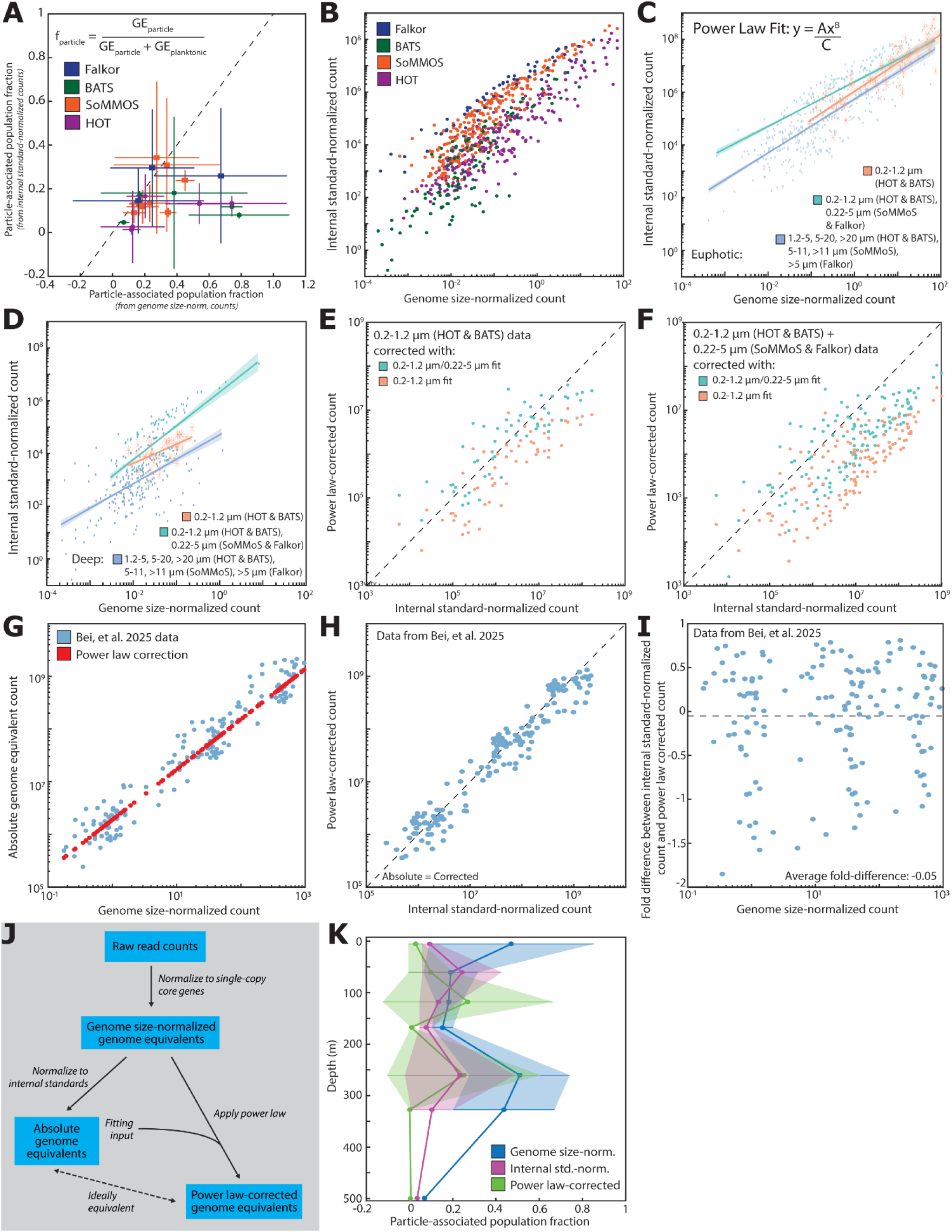
Relationship between genome size-normalized and internal standard-normalized genome equivalent counts for *Prochlorococcus* across size fractions. (A) Comparison of fraction of *Prochlorococcus* in particle-associated size fractions calculated using genome size-normalized or internal standard-normalized genome equivalents. (Note that for Falkor and SoMMoS cruises, the estimate is a lower bound on particle-associated fraction as the smallest size fraction is 0.22-5 μm.) (B) Relationship between genome size-normalized and internal standard-normalized *Prochlorococcus* genome equivalents across cruises shown in Fig. 1A. (C) Power law relationship between genome size-normalized and internal standard-normalized genome equivalents for euphotic (<= 150 m) data binned into 3 groups: free-living (0.2-1.2 μm from HOT and BATS), small (0.2-1.2 μm from HOT and BATS, 0.22-5 μm from Falkor and SoMMoS), and particle-associated (1.2-5 μm, 5-20 μm, and >20 μm from HOT and BATS, 5-11 μm, >11 μm from SoMMoS, and >5 μm from Falkor). (D) Power law relationship between genome size-normalized and internal standard-normalized genome equivalents for deep (>150 m) data with the same binning scheme as in C. (E) Comparison of free-living/euphotic (orange, average fold-difference −0.41) and small/euphotic (teal, average fold-difference −0.73) power law-correction on 0.2-1.2 μm data from HOT and BATS. (F) Comparison of free-living/euphotic (orange, average fold-difference 1.35) and small/euphotic (teal, average fold-difference 0.06) power law-correction on the combined 0.2-1.2 μm data from HOT and BATS and 0.22-5 μm from Falkor and SoMMoS. (G) Power law correction (small/euphotic) applied to independent dataset from Bei, *et al*. 2025 ^26^ (also using *Thermus* internal standard-normalized counts). (H) Comparison of absolute total *Prochlorococcus* genome equivalents reported by Bei, *et al*. 2025 normalized using the *Thermus* standard and the power law corrected *Prochlorococcus* genome equivalents calculated using the non-normalized data from Bei, *et al*. 2025. The dashed line represents a 1:1 equivalence of the quantities. (I) Comparison of the non-normalized data (genome size-normalized counts) and the fold difference between the internal standard-normalized and power law-corrected counts, showing an average fold difference of −0.05. (J) Schematic showing relationship between raw reads, genome-normalized counts, internal standard-normalized counts, and power law-corrected counts. (K) Comparison of fractions of *Prochlorococcus* calculated using genome size-normalized genome equivalents, internal standard-normalized genome equivalents, and power law-corrected genome equivalents.

However, we hypothesized that archival metagenomic data might still be useable if a general quantitative relationship exists between genome size- and internal standard-normalized counts. To test this, we investigated whether such a relationship holds across our samples. We compared the *Prochlorococcus* genome size-normalized genome equivalent counts to internal standard-normalized counts in the same samples across all size fractions from several different expeditions (Fig. 2B). While genome size-normalized counts were several orders of magnitude lower than internal standard-normalized counts, there was a similar relationship between the counts across the different datasets. When we plotted the genome size-normalized counts on a histogram, the distribution had a heavy tail and decayed roughly linearly across multiple orders of magnitude, suggesting a power law distribution. To test whether the data could accurately be fit by a power law, we sought to find the simplest binning strategy that would produce a fit that would also apply to each individual cruise sample set. Fitted power laws have the form *y* = *Ax^B^*, where y is the internal standard-normalized count, *x* is the genome size-normalized count, and *A* and *B* are fitting parameters. For each binning strategy, we tested the fitted power laws against the pooled dataset using a 100-fold cross validation with an 80/20 fit/test split (detailed example in Supp. Fig. 2A). Briefly, the pooled cruise data were split into bins, and in each bin, 80% of the data were used to fit a power law and the remaining 20% were used to test and quantify errors. This process was repeated 100 times. We calculated average fold-differences between the genome size-normalized and power law-corrected data (Supp. Fig. 2B), using these values as a third parameter for an adjusted power law of the form: 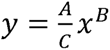, adding *C*, which is the average fold-difference determined from the cross validation. We used the average fold-difference, as well as the spread in power law-corrected values for a given internal standard-normalized count (Supp. Fig. 2C), to evaluate different binning strategies. Ultimately, the data were first split by depth (euphotic: <= 150 m, deep: >150 m) and then pooled in three ways based on filter size: 1) a “free-living” dataset of the 0.2-1.2 μm samples from HOT and BATS, 2) a “small size” dataset of the “free-living” dataset plus the 0.22-5 μm samples from Falkor and SoMMoS, and 3) a “particle-associated” dataset of the 1.2-5, 5-20, and >20 μm samples from HOT and BATS, 5-11, >11 μm samples from SoMMoS, and >5 μm samples from Falkor (Fig. 2C,D, Supp. Fig. 3E-J, Supp. Table 3). The resulting power laws quantitatively reproduced the relationship between genome size- and internal standard-normalized counts with an average fold-difference between the absolute and power law-corrected counts of <0.5 across all bins (except the “free-living/euphotic” bin, which is based on a small dataset and has an average fold-difference of 0.84). (See Supp. Fig. 3 for additional information on various binning strategies we tested.)

We also tested whether the broader “small” size bin could be used to correct samples from the free-living size fraction, given that we used a much larger dataset when fitting. Using only the 0.2-1.2 μm data from HOT and BATS, we applied both the free-living/euphotic and small/euphotic corrections (Fig. 2E, orange and teal respectively) and found a slight increase in the average fold-difference using the small/euphotic correction. Next, we compared the same corrections when applied to the combined 0.2-1.2 μm data from HOT and BATS and 0.22-5 μm data from SoMMoS and Falkor (Fig. 2F), where the fold-difference was significantly lower using the small/euphotic correction. These comparisons show that using the broader small/euphotic correction factor is more accurate if the cruise sampling scheme does not have a true free-living *Prochlorococcus* size fraction (i.e., 0.2-1.2 μm).

For a final validation of the power law correction, we tested this method using an independent metagenomic dataset that also used internal standards ^26^. Bei*, et al*. 2025 collected surface metagenomes, filtering the entire water sample onto a 0.22 μm filter, and extracted the samples with several types of internal genomic standards, including *Thermus thermophilus* ^26^. Using their data, we obtained *Prochlorococcus* genome size-normalized counts and corresponding internal standard-normalized counts in each sample (Fig. 2G, blue dots; see also Methods). Given the mixed size range in their metagenomes and based on our previous testing, we used our power law for the “small/euphotic” bin to calculate power law-corrected counts from their genome size-normalized counts (red dots). There was quantitative agreement between the data and our independently fit power law (Fig. 2H,I, average fold-difference of −0.05). Further improvement of our correction method awaits the availability of more size-fractionated marine metagenomic datasets with internal standards, but we are encouraged by our results thus far.

Given the limited sample size used to calculate the power laws based on data from the 0.2-1.2 μm size fraction, we focused on assessing optimal dataset sizes and error convergence with the four other fits (i.e. small/large, euphotic/deep). The average difference between internal standard-normalized and power law-corrected counts converges with about 50 metagenomic samples (Supp. Fig. 4A), and the percent error approaches 0 with approximately 100 samples (Supp. Fig. 4B). For robust use of this power law correction (schematic in Fig. 2J), we recommend using datasets with at least 50 metagenomic samples per bin. For the purposes of this pilot study, we aimed to use all available data, although of course the power law conversion would be more accurate with more samples from more specific size fractions.

Although the combined dataset from HOT, Falkor, BATS, and SoMMoS did not have 50 samples per depth/size bin, we tested how particle-associated fractions calculated from genome size-normalized counts, internal standard-normalized counts, and power law-corrected counts would compare across datasets. Given the limitations on our size fractionation and dataset sizes, we are presenting the lower bound on the *Prochlorococcus* particle-associated fraction, since these data average the particle-associated fractions from HOT and BATS with a true free-living size fraction (0.2-1.2 μm) and the lower-bound particle-associated fractions from SoMMoS and Falkor with a mixed small size fraction (0.22-5 μm), and we are using the “small/euphotic” correction bin to be consistent across cruises. We found that the population fractions calculated from power law-corrected counts were similar to the “true” fractions determined from internal standard-normalized counts (Fig. 2K, purple and green). Further, population fractions determined using genome size-normalized counts were not even able to capture the general shape of the data (Fig. 2K, blue), failing to capture an increase in particle-associated fraction between 0 and 75 m present in both internal standard-normalized and power law-corrected count data. Even with the smaller than ideal sample size, the power law correction was able to more closely match the depth profile determined from internal standard-normalized counts. Thus, the power law correction could be applied to metagenomic datasets extracted without internal standards to enable quantitative comparison between samples, such as calculating (lower bound) particle-associated fractions of *Prochlorococcus* populations or, more broadly, shifts in microbial community composition between free-living and particle-associated states.

### Trends in particle-associated populations from TARA Oceans data

The TARA Oceans Project collected size-fractionated samples for metagenomic analysis across the global oceans ^19–21^, providing an ideal test case for investigating *Prochlorococcus* particle association across different ocean regions by applying the power law correction. Since we were able to validate the “small/euphotic” power law correction with an independent dataset, we focused on euphotic samples from 0-150 m and binned the data as free-living (0.22-1.6 μm) or particle-associated (>1.6 μm), before applying the corresponding power law conversion (see Methods). Our final dataset comprised averaged particle-associated fractions across 47 locations collected between 0-80 m, excluding samples from the Arctic and Southern Oceans and only considering locations with a paired free-living and particle-associated sample (Fig. 3A).

**Figure 3.**
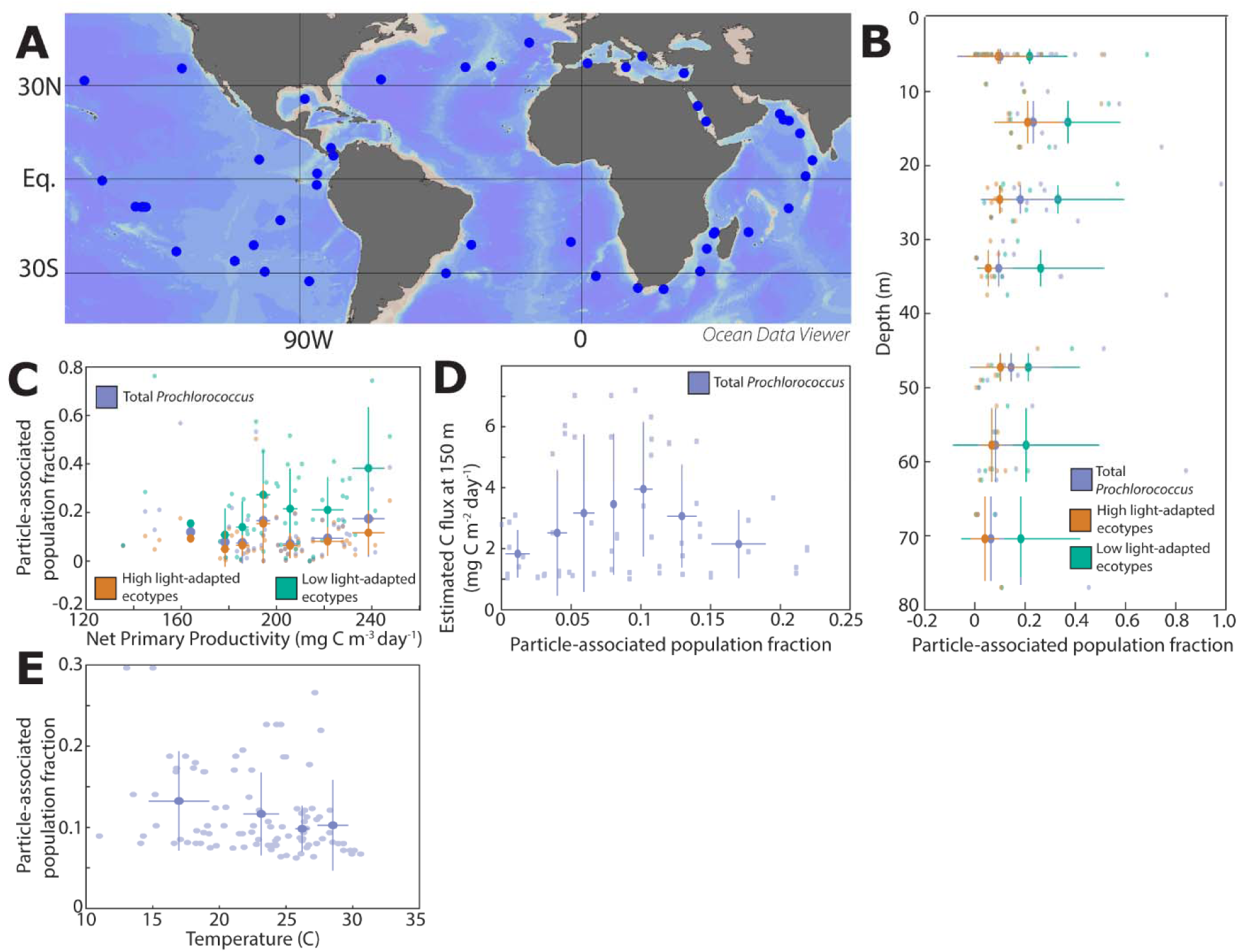
Power law-corrected particle-associated *Prochlorococcus* population fractions as a function of environmental parameters (data from TARA Oceans Project). (A) TARA sampling locations that passed quality control, detection limits, and had at least one free-living and particle-associated sample with identified *Prochlorococcus* genomes. The free-living size fraction is defined as 0.2-1.6 μm, and particle-associated size fractions are >1.6 μm. (B) Relationship between *Prochlorococcus* particle-associated population fraction and depth. Distributions for high light-adapted (orange), low light-adapted (green), and total population (purple) are shown. (C) Relationship between *Prochlorococcus* particle-associated population fraction and net primary productivity (high light: R=0.26, p=0.58; low light: R=0.81, p=0.03; total: R=0.36, p=0.42). (D) Relationship between *Prochlorococcus* particle-associated population fraction and estimated carbon export at 150 m (data from Guidi*, et al.* Nature 2016 ^31^). (E) Relationship between temperature and particle-associated *Prochlorococcus* fraction (R=-0.94, p=0.05).

We began by looking at depth distributions of particle-associated *Prochlorococcus* populations in the TARA data and asked whether these distributions were similar to the trends we observed from our internal standard-normalized metagenomic samples. Consistent with previously reported data ^5^ and our data reported in Supp. Fig. 1A-D, near the surface, low light-adapted ecotypes had higher particle-associated fractions (25.4±7.1%) and lower population size than high-light adapted ecotypes, while high light-adapted ecotypes had lower particle-associated fractions (9.1±5.7%) and a larger total population (Fig. 3B). Since high light-adapted *Prochlorococcus* cells are more abundant in these samples, the total particle-associated population fraction was similar to the high-light adapted fraction (12.3±6.5%). Overall, despite our data transformations, we found a similar average particle-associated fraction in the upper euphotic zone between our corrected TARA data and absolute data from HOT, BATS, Falkor, and SoMMoS.

We next set out to see if we could identify any ecosystem features that might influence the degree of particle-attachment among *Prochlorococcus* populations. We reasoned that populations could shift from free-living to particle-associated through several environmental mechanisms that would generally select for more or less particle association. Assuming that all cells have equal propensity for attachment, greater availability of attachment substrates or higher particle concentration would correlate with increased particle association. There could also be negative selection pressure exerted on free-living cells, such as through predation. Although there is insufficient data in the TARA metagenomes to determine specific mechanisms, we explored several hypotheses based on trends between environmental parameters and particle-associated *Prochlorococcus* fraction.

For the low-light adapted *Prochlorococcus* populations, the fraction of cells that were particle-associated increased with net primary productivity (Fig. 3C, green; R=0.81, p=0.03); this was not the case for high light-adapted population fractions (Fig. 3C, orange; R=0.26, p=0.58) (see Methods for details on correlations). Again, the total particle-associated population fraction closely matched the high light-adapted population and did not change with net primary productivity (Fig. 3C, purple; R=0.36, p=0.42), because these ecotypes dominate the population. Ocean regions with higher productivity have been linked to increased predation pressure on phytoplankton ^27^, which could more strongly impact free-living low-light ecotype cells of *Prochlorococcus*, which are larger than their high-light adapted counterparts and may be more attractive predator targets ^28,29^.

Increases in net primary productivity also result in increases in particulate organic carbon ^30^ and could be linked to increased particle sinking and carbon export, although this relationship is strongly affected by the ecosystem’s trophic relationships. Guidi, *et al*. 2016 reported estimates of carbon export at 150 m at TARA Oceans sampling sites, determined from particle imaging by an underwater vision profiler ^31^. Matching their carbon export measurements to *Prochlorococcus* particle association data obtained at the same TARA stations, we determined the relationship between particle association and carbon export: the two increased linearly until 10% of the *Prochlorococcus* population was particle-associated and decreased thereafter (Fig. 3D). The type of underwater vision profiler used in the TARA expedition is known to underestimate carbon export by more than two-fold due to the lack of detection of larger particles ^32^. Thus, if higher *Prochlorococcus* particle association is correlated with larger particles, then carbon flux could be underestimated at higher particle-associated fractions. Though we cannot test this directly with available data, we matched optical scattering data from water column samples to metagenomic samples at the same locations ^33^. Trends in the scattering parameters were consistent with particle size increasing as a function of both carbon export and particle-association fraction (for detailed methods and interpretation, see Supp. Fig. 6). These scattering patterns suggest that measured carbon export values at higher *Prochlorococcus* particle-associated fractions may be underestimated, indicating a potential linear relationship between particle association fraction and carbon export.

We also examined trends in particle association in relation to physical oceanographic conditions rather than productivity-related measures. The fraction of particle-associated *Prochlorococcus* decreased with temperature (R=-0.94, p=0.05, Fig. 3E) and increased with seawater density (R=0.82, p=0.05, Supp Fig. 5A). The fraction also increased with dissolved inorganic carbon (DIC) (R=0.87, p=0.03, Supp. Fig. 5B). These trends—high DIC, low temperature, and high density—are characteristic of upwelling ^34^ of deep water to the surface and align with our observations that a greater fraction of the *Prochlorococcus* population is particle-associated below the euphotic zone (Fig. 1D). Supporting this upwelling signal, DIC strongly correlated with nitrate and phosphate concentrations (Supp. Fig. 5C,D). However, neither nitrate nor phosphate showed a direct relationship with particle association (Supp. Fig. 5E,F), potentially because large differences in particle association between upwelled and euphotic water masses obscured any relationship within individual regions.

To test whether nutrient concentrations affect particle association independent of upwelling signals, we performed a residual analysis conditioning on DIC. After removing DIC-related variation, nitrate residuals positively correlated with particle association residuals (Supp. Fig. 5G), whereas phosphate residuals showed negligible correlation (Supp. Fig. 5H). These results suggest that particle association is primarily driven by DIC (as an upwelling proxy)—i.e. that the particles with attached *Prochlorococcus* had already sunk below the euphotic zone and were brought back up by upwelling. The correlation with nitrate but not phosphate is difficult to interpret other than to say that it is independent of upwelling dynamics.

We limited the analyses thus far to sampling locations where *Prochlorococcus* genomes were found in both the small and large size fractions, since it seems unlikely that *Prochlorococcus* would be present only on particles in a given water body. However, in all oceans except the Red Sea, there were multiple sites where *Prochlorococcus* genomes were exclusively identified in particle-associated samples (Indian Ocean: 8/22 samples, Mediterranean: 11/17, North Atlantic: 3/12, North Pacific: 7/13, Red: 0/4, South Atlantic: 5/11, South Pacific: 12/26). If we include these data in our analysis of the depth distribution of particle association, the estimated fractions dramatically increase to 40.6±10.3% for high light-adapted ecotypes, 50.1±8.8% for low light-adapted ecotypes, and 42.9±9.2% for the total population (Supp. Fig. 7A). Our analysis also excluded sampling locations from the Arctic or Southern Oceans, since *Prochlorococcus* is not generally found past 40 N/S ^1,4,25^. However, at all but one sampling location from these oceans, *Prochlorococcus* cells were identified only in large size fractions (Arctic 19/20, Southern 6/6).

As it is highly unlikely that *Prochlorococcus* is actively growing at these latitudes, we hypothesized that these genomes—as well as those from sampling sites in other oceans where *Prochlorococcus* was only identified in the large size fraction—were from particles that were widely distributed by ocean currents or by upwelling events. When we included the physical ocean parameters from these sampling sites and repeated our analysis relating particle association fraction to environmental parameters, we found that the correlation significance between seawater density or temperature and particle-associated fraction increased (R=0.79, p=0.03 and R=-0.99, p=0.01, respectively; Supp. Fig. 7B, C), suggesting that these locations may have sampled pockets of upwelled water with no free-living *Prochlorococcus*. One caveat is that none of these analyses reveal whether particle-associated (or free-living) *Prochlorococcus* are alive, however, these results may also indicate that we are underestimating the particle-associated *Prochlorococcus* population.

## Future Directions

This study has produced a scalable method for quantifying the fraction of particle-associated *Prochlorococcus* cells from metagenomic datasets that had been sequenced without an internal standard. Future work could apply these power law corrections to re-analyze archival ocean data and extend these conversions to other microbial groups of interest, although it is likely that individual microbial groups would require different correction factors.

The study was *post facto* not *ab initio*–i.e. designed to take advantage of existing resources for a pilot study. Two cruises—Falkor and SoMMoS—had already been collected and size fractionated, meaning we had no input on filter sizes. To maximize the potential of our approach in future expeditions investigating particle-associated *Prochlorococcus*, we propose the following sampling scheme: at each sampling site and depth, filter water using a 1.2 μm filter to capture particles and aggregates and a 0.2 μm filter to capture free-living cells. Simultaneously, a sample from the 0.2-1.2 μm size fraction should be stored for cell counting by flow cytometry. Bottle incubations of 1.2 μm filters used to trap particles with 0.2 μm-filtered, minimally-amended seawater would also provide opportunities to isolate living, particle-associated *Prochlorococcus* cells. This sampling scheme is built on *Prochlorococcus*’s cell size and intended to separate free-living and particle-associated cells. However, our delineation between “free-living” and “particle-associated” is empirically defined and will certainly be affected by sample collection effects, such as filter loading and sample volume ^35^. We offer it to begin to build an argument for a standardized size fractionation protocol that would eliminate some of the variability we encountered.

## Acknowledgements.

Thank you to the DeLong lab at the University of Hawai’i for Falkor expedition filters, the Sher Lab at the University of Haifa for SoMMoS expedition filters, and to the crew and marine technicians on the HOT346, BATS AE2412, SoMMoS, and FK180310 cruises for sample collection. The authors would also like to thank the TARA Oceans Consortium for open access to sampling schemes and datasets, Paul Berube for comments on the manuscript, Allison Coe and Nikolai Radzinski for assistance on cruise preparation, and the BioMicro Center at MIT for sequencing support.

## Author Contributions

M.A.-D. designed the study. A.J.S., S.M.S., S.J.B., and K.L. collected size-fractionated samples. M.A.-D., K.J., S.P., and K.C. extracted metagenomic samples. N.V. performed metagenomic analyses of field samples with input from M.A.-D. and J.M. M.A.-D. performed data processing and statistical analyses with help from N.V. M.A.-D., K.J., J.M., N.V., and K.C. were supervised by S.W.C. M.A.-D. wrote the paper with input from all authors.

## Methods

### Metagenome extraction

Serially-filtered samples were obtained from 3 ocean sampling expeditions: Falkor FK180310 in 2018 at Station ALOHA, SoMMoS (Southeastern Mediterranean Monthly cruise Series) in 2018-2019 at the Texas-Haifa Eastern Mediterranean Observatory 2 ^22^, and BATS AE2412 in 2024 from the Bermuda Atlantic Time-series Study site (see Supp. Table 1). We obtained the following size fractions: for FK180310, 0.22-5 μm and >5 μm; for SoMMoS, 0.22-5 μm, 5-11 μm, and >11 μm; for BATS AE2412, 0.2-1.2 μm, 1.2-5 μm, 5-20 μm, and >20 μm. DNA was extracted as described in ^5^ and modified from ^36,37^. DNA standards (*Thermus thermophilus* HB27, ATCC BAA-163D-S) were added according to estimated DNA yields at each depth. High-throughput libraries were prepared using Nextera FLEX (Illumina) and sequenced with 300 nucleotide paired-end reads using NextSeq 500 (Illumina). We also included previously reported data ^5^ collected at Station ALOHA (serially-filtered samples collected during HOT346 with size fractions 0.2-1.2 μm, 1.2-5 μm, 5-20 μm, and >20 μm) in our metagenome dataset.

### Taxonomic classification and quantification

To quantify high light-adapted and low light-adapted ecotypes, we classified metagenomics reads using ProSynTax, a protein dataset curated for taxonomic classification of *Prochlorococcus*, along with a modified version of the dataset’s accompanying workflow ^38^. Modifications to the workflow were made to enable the downloading and processing of TARA metagenomics samples from the NCBI’S SRA repository using the SRA Toolkit (RRID:SCR_024350), available through NCBI’s SRA repository (accessions provided in Supp. Table 4), as well as normalizing samples using internal standards. Briefly, samples were checked for quality using BBDuk v.39 ^39^, mapped to the internal standard reference genome (*Thermus thermophilus)*, and mapped reads were extracted using Seqtk v1.4 ^40^. To determine genome equivalents, we assumed that each *Prochlorococcus* cell has one copy of its chromosome ^41^ and used a set of 424 single copy core genes (SCCGs) from the CyCOG6 database ^42^ to identify and normalize read counts. Finally, to account for genome length variation across *Prochlorococcus* clades in taxonomically-classified reads, we aligned reads against the SCCG set, using DIAMOND v.2.1.11 ^42,43^. Genome-length-corrected values (called genome equivalents) were then obtained following:

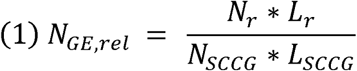

where *N_GE,rel_* is the number of genome equivalents (relative), *N_r_* is the number of reads mapped to *SCCGs, L_r_* is the length of the reads mapped to SCCGs in amino acids, *N_SCCG_* is the number of SCCGs, and *L_SCCG_* is the mean length of the SCCGs in amino acids.

### Sequencing efficiency and absolute genome equivalent calculation

To obtain absolute genome equivalents by normalizing to internal standards, we quantified sequencing efficiency by dividing standard molecules added by standard molecules recovered (Eqs. 2, 3).

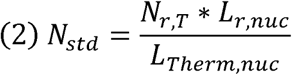

where *N_std_* is the number of standard molecules recovered, *N_r,T_* is the number of reads mapped to the *Thermus* genome, *L_r,nuc_* is the read length in nucleotides, and *L_Therm,nuc_* is the length of the *Thermus* genome in nucleotides.

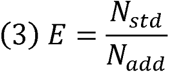

where *E* is the efficiency, *N_std_* is the number of standard molecules recovered, and *N_add_* is the number of standard molecules added.

Finally, we divided the genome equivalents results from the ProSynTax workflow by the sequencing efficiency to obtain absolute genome equivalents (Eq. 4).

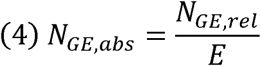

where *E* is the efficiency, *N_GE,rel_* is the number of relative genome equivalents, and *N_GE,abs_* is the number of absolute genome equivalents.

### Particle-associated population fraction calculation

We calculated the particle-associated fraction of the total *Prochlorococcus* population, *f_particle_*, by dividing the sum of the number of genome equivalents in the particle-associated size fractions (ΣGE_particle_) by the sum of the total number of genome equivalents in both the free-living size fraction and the particle-associated size fraction (ΣGE_particle_ + ΣGE_free-living_):

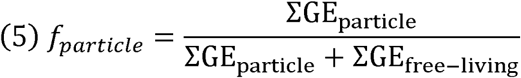

The explicit summations for the particle-associated fraction calculated from size fraction genome equivalents for each cruise are listed below (Eq. 6 for HOT, Eq. 7 for BATS, Eq. 8 for SoMMoS, and Eq. 9 for Falkor):

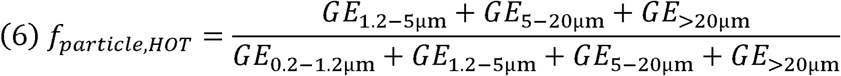

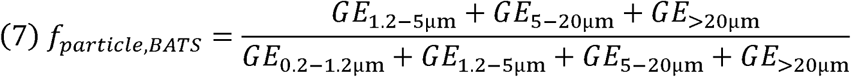

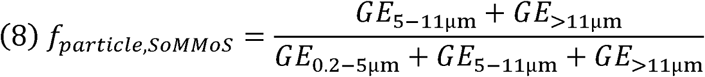

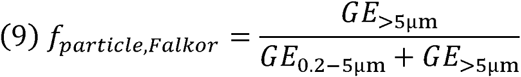

### Population size estimates

For flow cytometry counts related to the SoMMoS cruises, we use the counts from the closest corresponding date and depth from Reich, *et al.* 2022 ^22^. For counts related to the FK180310 cruise, we used the reported core water column data from Wilson and Poulos 2018 ^44^. Finally, for counts related to HOT346 and BATS AE2412, we averaged depth profiles of *Prochlorococcus* concentrations from 2020-2022 (see Supp. Table 2 for public repository information; for HOT, data were obtained via the Hawai’i Ocean Time-series HOT-DOGS application; University of Hawai’i at Mānoa, NSF Award #2241005). At each depth with a paired metagenomic and flow cytometry measurement, we calculated the total *Prochlorococcus* population as the free-living cell count divided by the free-living fraction. Finally, given low resolution in the data, we averaged the free-living and particle-associated populations between 0-150 m at each cruise location to get an average count in the euphotic zone (see full data sources, aggregation, and averaging in Supplementary Table 2).

### Power law validation

To validate the power law correction for “small/euphotic” samples (i.e. the power law fitted on data from 0.2-1.2 μm size fractions from HOT and BATS and 0.22-5 μm from Falkor and SoMMoS), we re-analyzed an independent metagenomic dataset with internal standards ^26^. Bei*, et al*. 2025 collected surface metagenomes, filtering the entire water sample onto a 0.22 μm filter, and extracting the samples with several types of internal genomic standards, including *Thermus thermophilus* ^26^. Their metagenomic samples were processed using the workflow detailed above (“Taxonomic classification and quantification” and “Sequencing efficiency and absolute genome equivalent calculation”) with identical parameters to other analyses in this paper to obtain absolute genome equivalent counts for *Prochlorococcus*. To assess the accuracy of power laws, we calculated absolute errors (defined as the difference between the true read count and the corrected read count) and fold-difference errors (defined as the absolute error divided by the true read count). We compared fold-difference distributions for the total dataset and individual cruises to ensure that there were not large differences in error distributions. P-values for comparisons between the fold-differences between the corrected data using the entire dataset or individual cruise datasets were calculated using 2-tailed t-tests (analysis performed in MATLAB 2023b).

### TARA Oceans Project particle-associated fraction estimation

To estimate particle-associated populations using data from the TARA Oceans Project ^19–21^, we first classified metagenomic reads using the ProSynTax pipeline described above (for sample accession numbers, see Supp. Table 4), beginning with 1799 metagenomic samples, of which 1170 passed initial quality control and detection limit thresholding. Next, we matched samples with environmental metadata ^33,45^. (Note that in the TARA carbonate chemistry dataset ^33^, dissolved inorganic carbon is reported as “total carbon”.) Next, we applied the “small/euphotic” power-law conversion to size-and depth-binned samples, resulting in power law-corrected genome equivalents for 218 free-living samples and 411 particle-associated samples. Particle-associated fractions were calculated as follows: for all samples at a given latitude/longitude, samples were either classified as “free-living” (0.2 μm-1.6 μm), “particle-associated” (>1.6 μm), or “other” (size fractions spanning free-living and particle-associated sizes or samples without size fractionation). Samples in the “other” category were discarded from the dataset. If the dataset contained exact replicates (i.e., samples collected at the same latitude/longitude with the same collection and pre-filter sizes), the genome counts were averaged. If a given location had at least 1 sample in both the free-living and particle-associated fractions and *Prochlorococcus* genomes were identified in both size fractions, the data were kept. Finally, particle-associated fractions were calculated as described above in “particle-associated population fraction calculation”.

### Correlation analysis for TARA data

Correlation coefficients were calculated using pairwise linear correlation, and reported p-values are testing whether the correlation is significantly different from 0 (2-tailed correlation for Pearson’s correlation using a student’s t-distribution). We note that due to the noise in the data, exact p-values and correlation coefficients are sensitive to data binning, hence we only discuss general trends. Analysis was performed in MATLAB 2023b.

## Data Availability

The raw metagenomics data generated for this study are available through the NCBI SRA with accession code PRJNA1512989 with associated metadata provided in Supplementary Table 1. The NCBI SRA accession codes for all TARA samples analyzed in this study are available in Supplementary Table 4.

