## Supplementary Information for "The distribution of particle-associated *Prochlorococcus* across the global oceans"

**Supplementary Information for “The distribution of particle-associated *Prochlorococcus* across the global oceans”, Anjur-Dietrich, *et al.***

**Supplementary Table 1. Metadata for all metagenomic data generated for this paper**, including sample name, original cruise, depth, filter size, water volume filtered, latitude and longitude, and date sample collected. Provided as an Excel sheet.

**Supplementary Table 2. Flow cytometry cell count data and calculations of *Prochlorococcus* population sizes.** For each cruise, average flow cytometry counts with depth are listed (data sources specified). Particle-associated cell counts are estimated based on population fractions from metagenomic data and counts from flow cytometry. Provided as an Excel sheet.

Note: Quantitative metagenomic analyses allow determination of the genome equivalent count of free-living and particle-associated *Prochlorococcus* and thus the fraction of the population in either state. Analysis by flow cytometry directly yields the number of free-living cells. However, neither measure can directly count the number of particle-associated cells. Although flow cytometry and metagenomic samples were not paired across the four cruises, we estimated the population size of particle-associated cells (see also, Supp. Fig. 1E, Methods). We also compared absolute metagenomic counts (genome equivalents mL^-1^) with flow cytometry counts (cells mL^-1^) of free-living cells and found some variability across cruises, although generally the two measurements were on the same order of magnitude (see also, Supp. Fig. 1F). In previous studies, this discrepancy was partly attributed to *Prochlorococcus* cells in surface waters having lower chlorophyll (hence less red fluorescence) and falling below the limit of detection by flow cytometry ^1,2^; differences between our measurements may also stem from the lack of direct pairing between measurements. Overall, robust estimates of particle-associated population size require simultaneously collected flow cytometry and metagenomic samples.

**Supplementary Table 3. Parameters from power law fitting for various dataset binning strategies.** For each approach, the data is fit by $y=\frac{Ax^{B}}{C}$, where x is the genome size-normalized genome equivalent count, y is the corrected count, A is the y-intercept on a log-log scale, B is the slope on a log-log scale, and C is the average fold-difference from the 100-fold cross validation (see Methods and Supp. Fig. 2 for more information). Provided as an Excel sheet. (See also Supp. Fig. 3 for graphs of fitting for binning strategies)

**Supplementary Table 4. Accession numbers for all TARA samples used in this paper.** Provided as an Excel sheet.

**Supplementary Figure 1. Ecotype distribution of particle-associated fractions for the cruises described in Fig. 1.** (A-D) Fractions of *Prochlorococcus* population found in the particle-associated size fraction for BATS AE2412 (1.2-5, 5-20, >20 μm), HOT 346 (1.2-5, 5-20, >20 μm), SoMMOS (5-11, >11 μm), and FK180310 (>11 μm) cruises (A-D, respectively) with high light-, low light-adapted, and unclassified *Prochlorococcus* populations shown, as well as total *Prochlorococcus*. For SoMMOS and FK180310, the data presented are a lower-bound estimate of the particle-associated population fraction, as the smallest metagenomic size fraction spans 0.22-5 μm. (Note that for readability, we refer to HOT346 as “HOT”, BATS AE2412 as “BATS”, FK180310 as “Falkor”, and SoMMoS as “SoMMoS” throughout.) (E) Estimated population counts of particle-associated *Prochlorococcus* based on combining metagenomic data and flow cytometry. Flow cytometry and metagenomic samples were not directly paired across the four datasets, and in two cruises, the smallest size fraction likely contained small particles and aggregates. To obtain a best estimate, we matched flow cytometry counts to the closest corresponding date and depth from each cruise, then averaged *Prochlorococcus* concentrations between 0–150 m to determine particle-associated populations at each location (see Methods, Supp. Table 2). Then, we combined the fraction of *Prochlorococcus* genomes in the free-living size fraction (calculated from metagenomics data) and the number of free-living cells (calculated from flow cytometry data) to determine the total number of *Prochlorococcus* cells (total = free-living cell count / free-living population fraction). Then, we determined the number of particle-associated cells by multiplying the total cell count by the particle-associated fraction. Ideally, this calculation would be based on metagenomic and flow cytometry analysis of aliquots taken from the same seawater sample, meaning that our population estimates are rough given the indirect pairing. (F) Comparison of *Prochlorococcus* absolute genome equivalent counts in the smallest size fraction (0.2-1.2 μm for BATS AE2412 and HOT 346; 0.22-5 μm for SoMMoS and FK180310) determined by metagenomes (genomes mL^-1^) and flow cytometry (cells mL^-1^) for each cruise (see also Supp. Table 2).

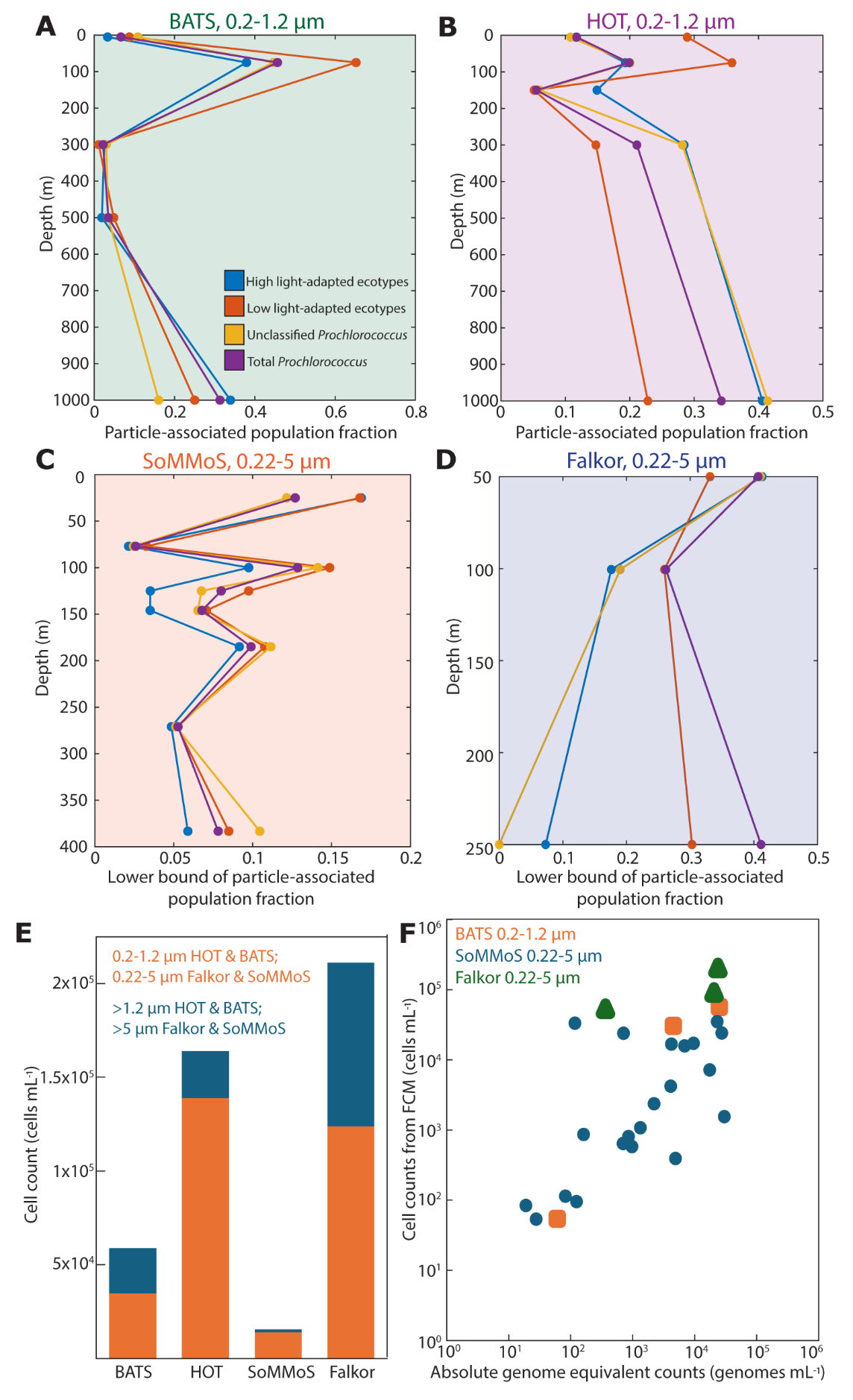

**Supplementary Figure 2. Fit/test cross validation example for four bins of euphotic/small, euphotic/large, deep/small, and deep/large (small: 0.2-1.2 μm from HOT and BATS, 0.22-5 μm from SoMMoS and Falkor; large: 1.2-5, 5-20, and >20 μm from HOT and BATS, 5-11, and >11 μm from SoMMoS, >5 μm from Falkor; euphotic: <=150 m; deep: >150 m).** (A) Schematic of 100-fold cross validation of power law fit using an 80%/20% fit/test split for each data bin. (B) Fold difference between the internal standard-normalized genome equivalents and the power law corrected genome-size normalized genome equivalents for each data bin (i-iv). (C) Comparison between the internal standard-normalized genome equivalents and the power law corrected genome-size normalized genome equivalents for each data bin with red dashed lines showing a 1:1 correspondence (i-iv).

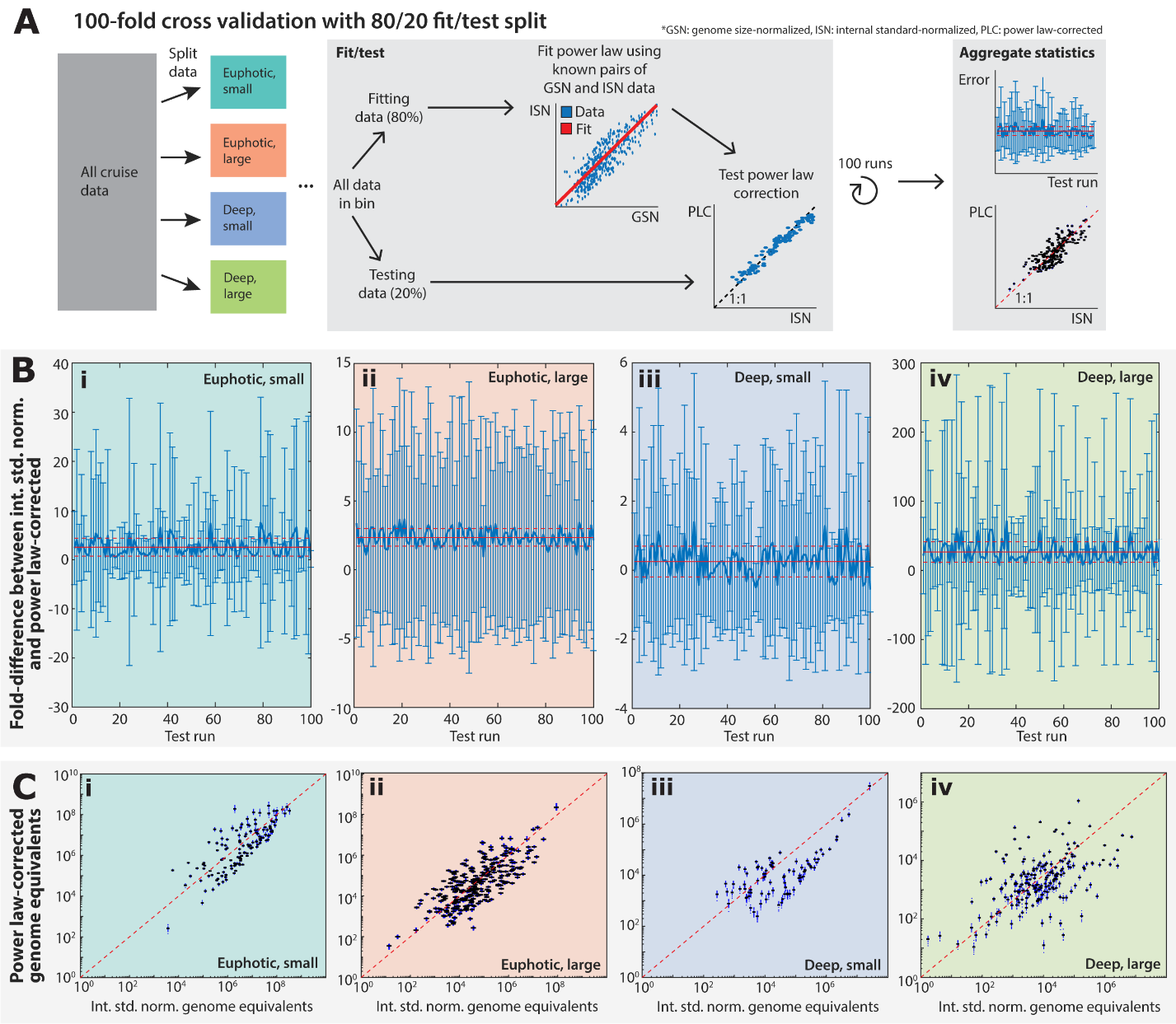

**Supplementary Figure 3. Assessment of various dataset binning strategies and associated power law fits. (For all fitting parameters, see Supp. Table 3)** (A) Raw Falkor dataset including 6 outliers is shown. These outliers were removed when aggregating the datasets for this paper. No other data were excluded. (B) Power law fitting for the entire aggregated dataset (HOT346, FK180310, BATS AE2412, SoMMoS) with the raw data in blue and fit in red with a shaded 95% confidence interval. (C) Power law fitting for data split between euphotic samples (depth <= 150 m, light blue points with red fit) and deep samples (depth >150 m, dark blue points with purple fit). (D) Power law fitting for data split between small size fraction samples (0.2-1.2 μm from HOT and BATS and 0.22-5 μm from SoMMoS and Falkor, light blue points with red fit) and large size fraction samples (1.2-5, 5-20, and >20 μm from HOT and BATS, 5-11, and >11 μm from SoMMoS, or >5 μm from Falkor), dark blue points with purple fit). (E) Power law fit for free-living, euphotic samples (0.2-1.2 μm for BATS and HOT only, depth <= 150 m). (F) Power law fit for small, euphotic samples (0.2-1.2 μm for BATS and HOT and 0.22-5 μm for Falkor and SoMMoS, depth <= 150 m). (G) Power law fit for particle-associated, euphotic samples (1.2-5, 5-20, >20 μm for BATS and HOT, 5-11, >11 μm for SoMMoS, and >5 μm for Falkor, depth <= 150 m). (H) Power law fit for free-living, euphotic samples (0.2-1.2 μm for BATS and HOT only, depth > 150 m). (I) Power law fit for small, deep samples (0.2-1.2 μm for BATS and HOT and 0.22-5 μm for Falkor and SoMMoS, depth <= 150 m). (J) Power law fit for particle-associated, deep samples (1.2-5, 5-20, >20 μm for BATS and HOT, 5-11, >11 μm for SoMMoS, and >5 μm for Falkor, depth > 150 m). (See also Fig. 2 for panels F-J)

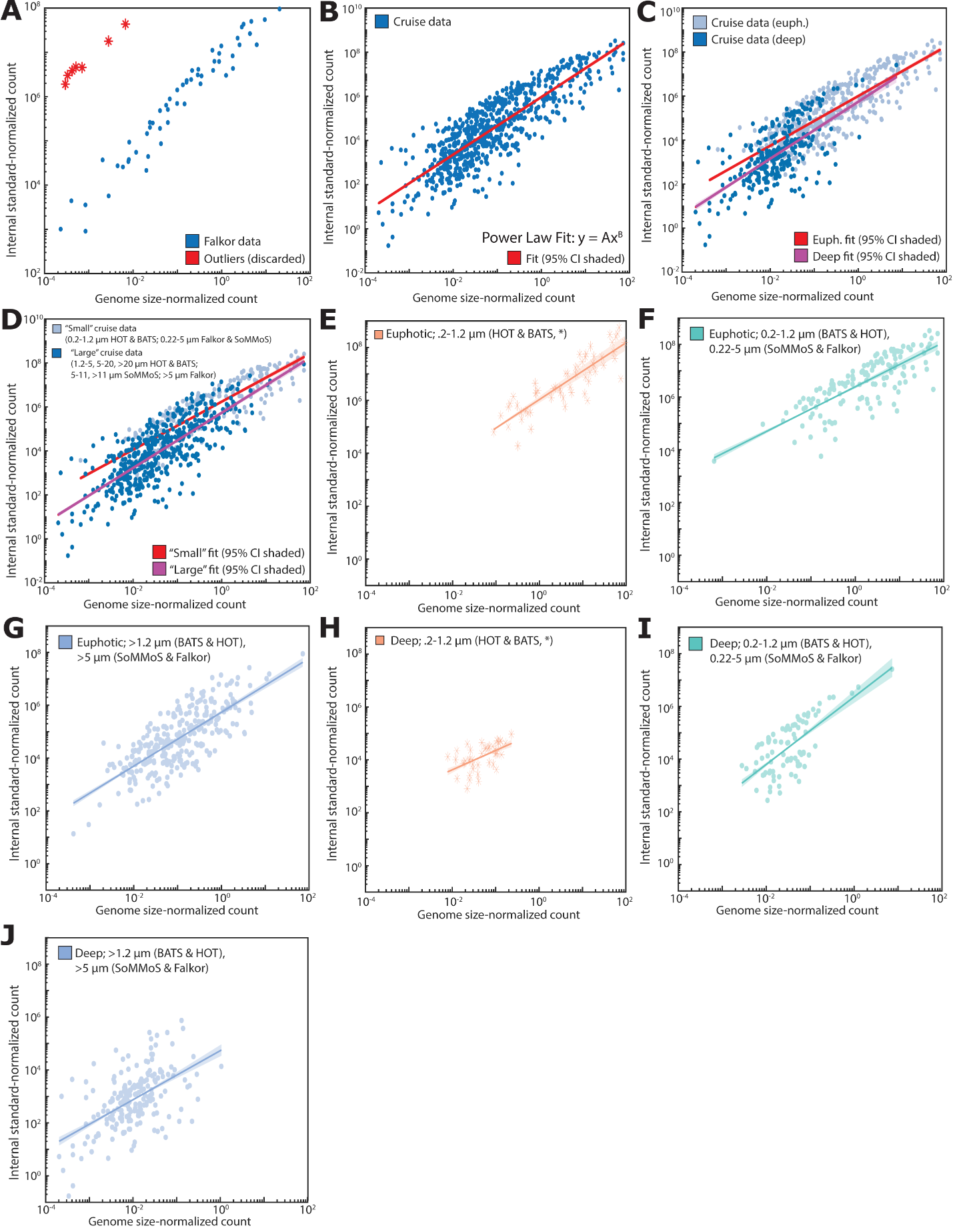
**Supplementary Figure 4. Error and convergence testing of power law corrections.** (A) Average difference between internal standard-normalized genome equivalents and power law-corrected genome equivalents as a function of increasing dataset size. (B) Average percentage error (std. dev./mean) on difference between internal standard-normalized genome equivalents and power paw-corrected genome equivalents as a function of increasing dataset size.

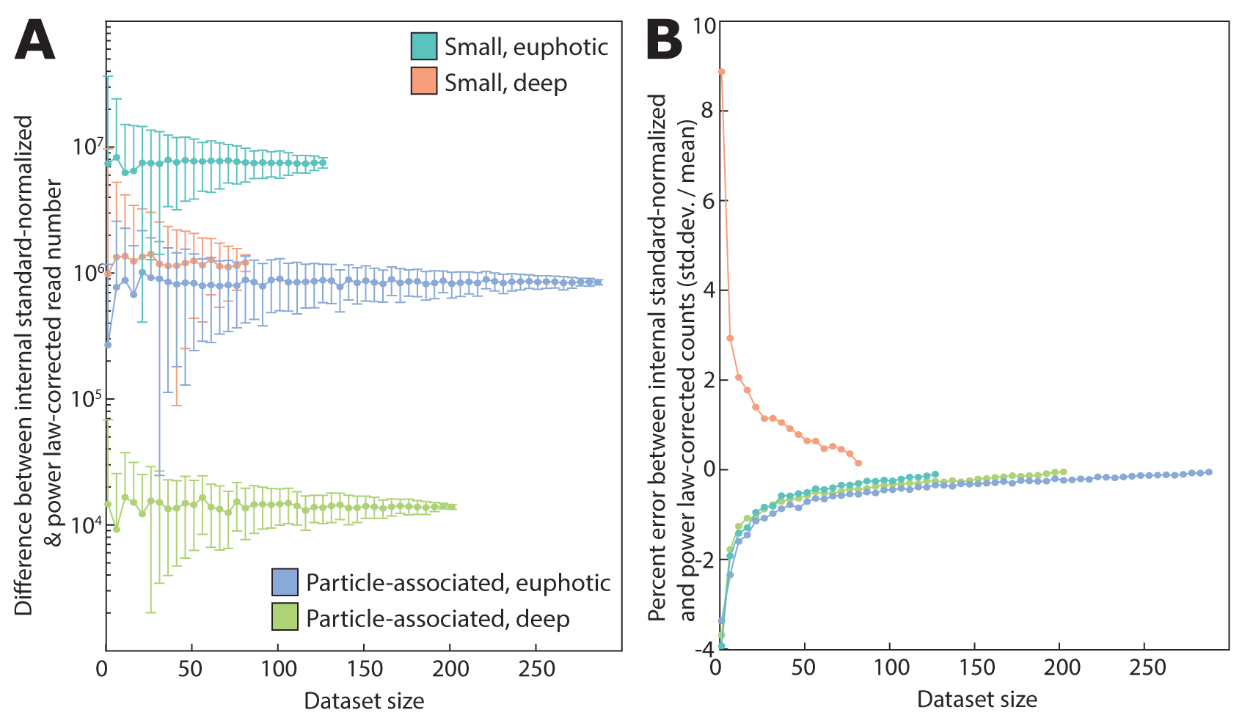

**Supplementary Figure 5. Additional analyses relating power law-corrected particle-associated *Prochlorococcus* population fractions to environmental parameters (data from TARA Oceans Project).** (A) Relationship between total *Prochlorococcus* particle-associated population fraction and seawater density (R=0.82, p=0.05). (B) Relationship between dissolved inorganic carbon (DIC) concentration and particle-associated fraction for high light-adapted (orange), low light-adapted (green), and total (purple) *Prochlorococcus* (total population: R=0.87, p=0.03). (C) Relationship between DIC and nitrate concentration in seawater samples (R=0.90, p=0.04). (D) Relationship between DIC and phosphate concentration (R=0.99, p<0.001). (E) Relationship between *Prochlorococcus* particle-associated population fraction and nitrate concentration (R=0.44, p=0.24). (F) Relationship between *Prochlorococcus* particle-associated population fraction and phosphate concentration (R=0.35, p=0.33). (G) Relationship between the residuals of nitrate concentration regressed on DIC and the residuals of particle-associated *Prochlorococcus* fraction regressed on DIC (R=0.76, p=0.14). (H) Relationship between the residuals of phosphate concentration regressed on DIC and the residuals of particle-associated *Prochlorococcus* fraction regressed on DIC (R=-0.31, p=0.62).

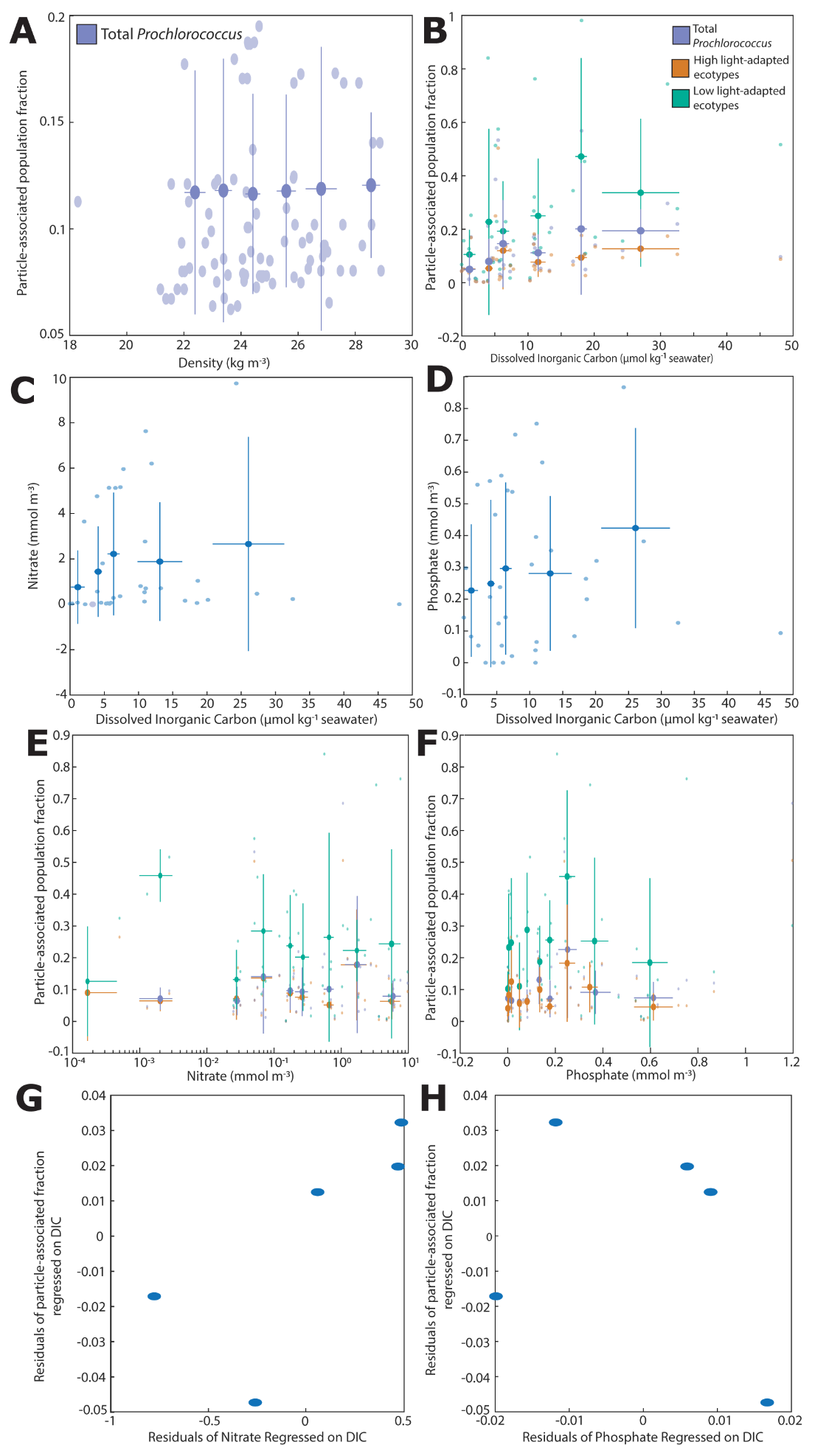

**Supplementary Figure 6. Additional data relating power law-corrected particle-associated *Prochlorococcus* population fractions to carbon flux measurements (data from TARA Oceans Project and Guidi*, et al.* 2016 ^3^).** (A) Relationship between particle backscattering coefficient (m^-1^) and estimated carbon export at 150 m (mg C m^-2^ day^-1^). (B) Relationship between beam attenuation coefficient (m^-1^) and estimated carbon export at 150 m (mg C m^-2^ day^-1^). (C) Relationship between the ratio of backscattering coefficient to beam attenuation coefficient (unitless) and estimated carbon export at 150 m (mg C m^-2^ day^-1^). (D) Relationship between *Prochlorococcus* particle-associated fraction and particle backscattering coefficient (m^-1^). (E) Relationship between *Prochlorococcus* particle-associated fraction and beam attenuation coefficient (m^-1^). (F) Relationship between *Prochlorococcus* particle-associated fraction and the ratio of backscattering coefficient to beam attenuation coefficient (unitless).

Note, we make the following assumptions in our interpretation of patterns in scattering coefficients: the samples had similar, heterogenous material composition with broad, non-specific absorption bands; fluorescence effects are neglected (see, for example, assumptions about “gelbstoff” and/or “detritus” in Dierssen 2010 ^4^ or Werdell, *et al.* 2018 ^5^). We looked at 1) backscattering coefficient, 2) beam attenuation coefficient (as a proxy for total scattering), and 3) the ratio of the backscattering coefficient to the beam attenuation coefficient (as a proxy for the backscattering ratio). If either estimated carbon flux or particle-associated population fraction are related to larger particles, then as these quantities increase, 1) backscattering should increase, 2) total scattering should increase, and 3) the backscattering ratio should decrease.

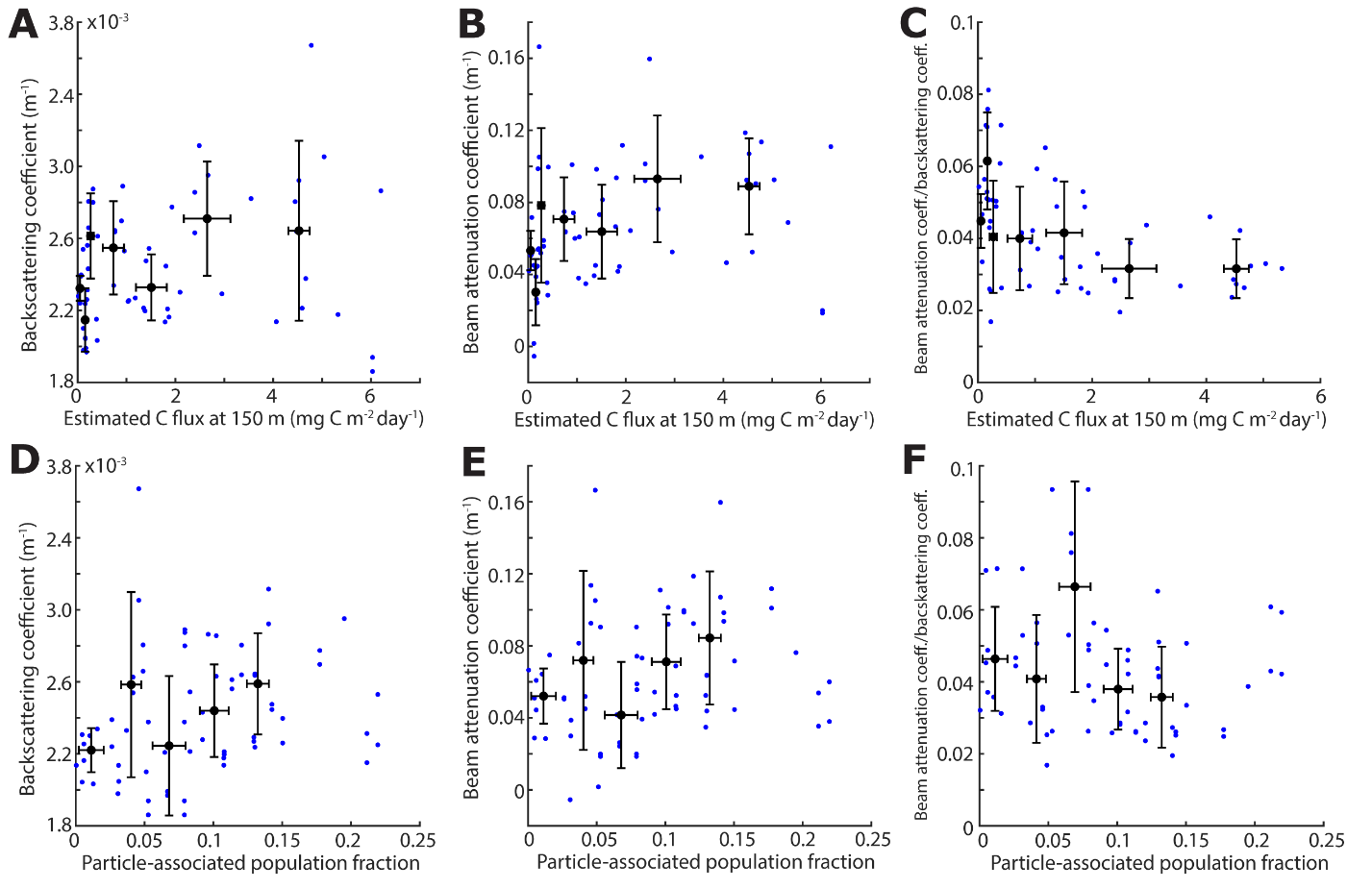

**Supplementary Figure 7. Additional plots for relating power law-corrected particle-associated *Prochlorococcus* population fractions to environmental parameters (data from TARA Oceans Project) including sampling sites where *Prochlorococcus* was only identified in large size fractions.** (A) Relationship between *Prochlorococcus* particle-associated population fraction and depth. Distributions for high light-adapted (orange), low light-adapted (green), and total population (purple) are shown. (B) Relationship between total *Prochlorococcus* particle-associated population fraction and seawater density (R=0.79, p=0.03). (C) Relationship between *Prochlorococcus* particle-associated population fraction (total population) and seawater temperature (R=-0.99, p=0.01).

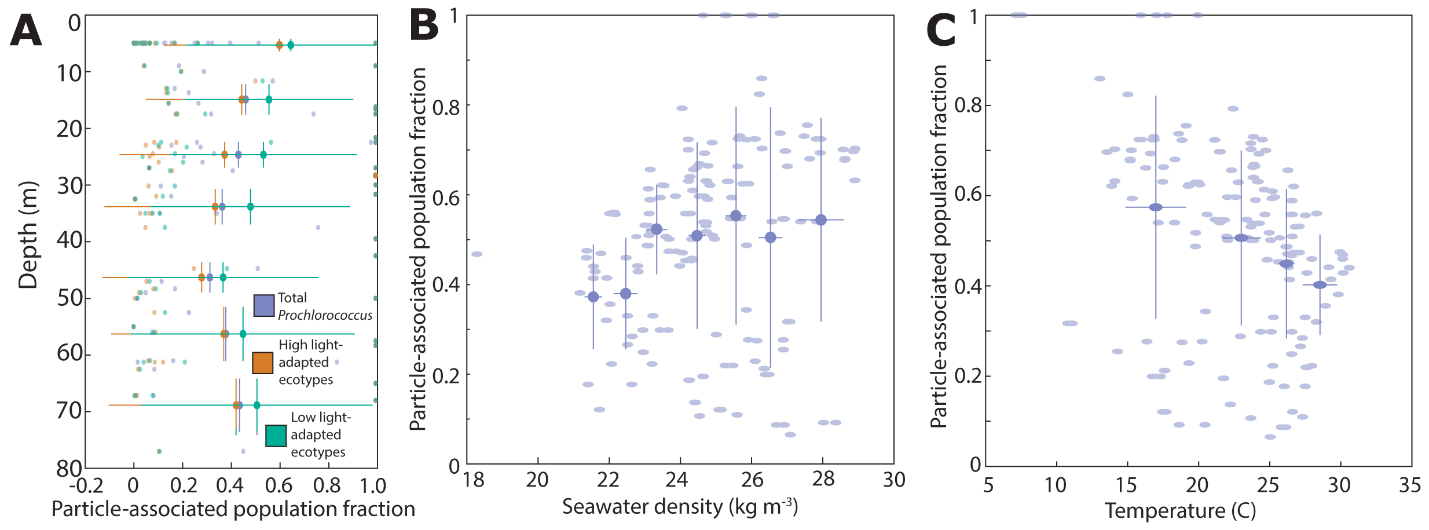
